# Aalbo1200: global genetic differentiation and variability of the mosquito *Aedes albopictus*

**DOI:** 10.1101/2023.11.21.568070

**Authors:** Jacob E. Crawford, Nigel Beebe, Mariangela Bonizzoni, Beniamino Caputo, Brendan H. Carter, Chun-Hong Chen, Luciano V. Cosme, Carlo Maria De Marco, Alessandra della Torre, Elizabet Estallo, Xiang Guo, Wei-Liang Liu, Kevin Maringer, Jimmy Mains, Andrew Maynard, Motoyoshi Mogi, Todd Livdahl, Noah Rose, Patricia Y. Scarafia, Dave Severson, Marina Stein, Sinnathamby N. Surendran, Nobuko Tuno, Isra Wahid, Xiaoming Wang, Jiannong Xu, Guiyan Yan, Don Yee, Peter A. Armbruster, Adalgisa Coccone, Bradley J. White

## Abstract

The mosquito *Aedes albopictus* transmits human viruses including dengue and chikungunya and is an extremely successful invasive species expanding into new regions of the world. New tools are needed to complement existing tools to help monitor and control this species. Genomic resources are improving for this species including genome reference sequences, and whole genome sequencing data will help to catalog genetic diversity in this species and further enable genetic analysis. We collected populations of *Ae. albopictus* from throughout its distribution and generated whole genome sequencing data from population samples. These data will be used to address a number of basic and applied questions for this species. Here, we show genetic differentiation patterns among the tropical and temperate forms, as well as finer scale genetic clustering at the regional and population scale. These data and results will be a valuable resource for further study and tool development for this species.

## Introduction

Human viruses transmitted by mosquitoes in the genus *Aedes* including dengue and chikungunya sicken over 100 million people every year throughout the tropical and subtropical parts of the world [1,2]. *Aedes albopictus* is native to South East Asia, but it is a remarkably successful invasive species that has expanded out of that native range into many temperate regions [3]. Most recently, *Ae. albopictus* has spread and established in a number of European countries, leading to an increase in local transmission of dengue in Mediterranean countries in recent years [4,5]. The dramatic success of this invasive species is due in part to its ability to overwinter in temperate regions facilitated by an ability to undergo a photoperiodic diapause [6–8], but climate change continues to increase the range of suitable habitat for both diapausing and non-diapausing populations of this species [9].

Without widely accessible vaccines against dengue and chikungunya [10], mosquito control is the primary tool in the effort to limit the public health burden of these diseases. Source control is laborious, expensive, and error prone as *Ae. albopictus* is a container breeder capable of exploiting very small and ephemeral water sources. Common chemical insecticides are losing efficacy against this species in some regions where levels of resistance are increasing [11,12]. Development of new tools to control and monitor this species is urgently needed.

Understanding the genomic landscape of genetic variability and global distribution of genetic diversity and differentiation is an important piece of the foundational knowledge in the effort to build new tools. Towards that goal, we assembled population samples from throughout the global distribution of *Ae. albopictus* and generated whole genome sequencing data to form a global genomes database we call Aalbo1200. In addition to our initial population genetic results of the dataset, we hope the underlying data will serve as a useful resource to help develop new tools and further our understanding of this mosquito species.

## Methods

### Mosquito samples

Mosquitoes were collected either as eggs using ovitraps, as larvae collected in small numbers from larval habitats, or as adults. Location details, sampling notes, and sample sizes are listed in Table 1. Samples were shipped to Verily for sequencing either as previously extracted DNA or as preserved larval or adult carcasses.

**Table 1.** Mosquito Population Samples. Sampling location (latitude and longitude) are approximate and location refers to the city or region where the sample was collected.

| Location | Country | Number individuals | Latitude | Longitude |
| --- | --- | --- | --- | --- |
| Salem | USA | 20 | 39.9771201 | -74.553223 |
| Manassas | USA | 20 | 38.4277735 | -78.288574 |
| New Zion | USA | 19 | 33.8443276 | -80.029241 |
| Jacksonville | USA | 20 | 30.1641263 | -81.804199 |
| Oak Hill | USA | 24 | 28.8644338 | -80.854497 |
| Vero_Beach | USA | 20 | 27.6386434 | -80.397274 |
| Pahokee | USA | 20 | 26.8200607 | -80.665335 |
| Palm Beach | USA | 20 | 26.9416595 | -80.573731 |
| New Orleans | USA | 20 | 29.9510658 | -90.071532 |
| Springfield | USA | 20 | 37.2298895 | -93.310451 |
| Houston 2001 | USA | 21 | 29.7934607 | -95.32832 |
| Houston 2018 | USA | 19 | 29.7934607 | -95.32832 |
| Hattiesburg | USA | 19 | 31.3271189 | -89.290339 |
| Lexington | USA | 20 | 38.0405837 | -84.503716 |
| Los Angeles | USA | 20 | 34.0522342 | -118.24368 |
| Oahu | USA | 10 | 21.4289349 | -158.00474 |
| Jardin Panteon | Mexico | 24 | 18.9967918 | -98.2289 |
| St. Augustine | Trinidad | 18 | 10.6473412 | -61.399777 |
| Cali | Colombia | 8 | 3.5828111 | -76.462458 |
| Barras das Carças | Brazil | 20 | -15.89162 | -52.261883 |
| Cachoiero de Itapemirim | Brazil | 20 | -20.846705 | -41.12022 |
| Gravataí | Brazil | 19 | -29.942289 | -50.990788 |
| Maceio | Brazil | 20 | -9.6453708 | -35.710982 |
| Manaus | Brazil | 20 | -3.1190275 | -60.021731 |
| Puerto Iguazú Misiones | Argentina | 15 | -25.597164 | -54.578599 |
| Sendai | Japan | 19 | 38.4277735 | 140.822754 |
| Wajima | Japan | 24 | 37.3905901 | 136.899196 |
| Tokyo | Japan | 20 | 35.8356284 | 139.746094 |
| Tanegashima | Japan | 25 | 30.609558 | 130.978878 |
| Okinawa | Japan | 9 | 26.2124013 | 127.680932 |
| Tainan | Taiwan | 16 | 24.126702 | 120.717773 |
| Beijing | China | 8 | 39.9771201 | 116.367188 |
| Shanghai | China | 17 | 31.3536369 | 121.376953 |
| Guangzhou | China | 20 | 23.1605633 | 113.137207 |
| Hainan | China | 20 | 19.3940679 | 109.643555 |
| Yunnan | China | 20 | 24.4271453 | 101.403809 |
| Chanthaburi | Thailand | 18 | 13.2029586 | 102.209022 |
| Ipoh | Malaysia | 24 | 4.597479 | 101.090106 |
| Kuala_Lumpur | Malaysia | 19 | 3.2721456 | 101.645508 |
| Jakarta | Indonesia | 24 | -6.17511 | 106.86504 |
| Sulawesi Urban | Indonesia | 13 | -2.1747706 | 121.223145 |
| Sulawesi Forest | Indonesia | 12 | -2.1747706 | 121.223145 |
| Sumba | Indonesia | 24 | -9.6993439 | 119.974053 |
| Papua | Indonesia | 12 | -4.4553223 | 137.136213 |
| Gelephu | Bhutan | 4 | 26.8725114 | 90.4927134 |
| Kathmandu | Nepal | 12 | 27.7172453 | 85.3239605 |
| Kunfunadhoo | Republic of Maldives | 8 | 5.1122262 | 73.0781579 |
| Jaffna | Sri Lanka | 7 | 9.6614981 | 80.0255465 |
| Torres Strait Islands | Australia | 60 | -10.05082 | 143.069406 |
| Honiara | Solomon Islands | 16 | -9.4343868 | 159.960989 |
| Limbe City | Cameroon | 20 | 4.0224656 | 9.1954432 |
| Awka | Nigeria | 20 | 6.2220043 | 7.0821162 |
| Morondava | Madagascar | 19 | -20.28401 | 44.3474121 |
| Tampon | La Reunion | 24 | -21.286278 | 55.519603 |
| Montpelier | France | 21 | 43.6134596 | 3.8864136 |
| Imperia | Italy | 20 | 43.8896857 | 8.039517 |
| Rome | Italy | 20 | 41.9027008 | 12.4962352 |
| Tirana | Albania | 20 | 41.3275459 | 19.8186982 |
| Krasnodar | Russia | 14 | 45.036035 | 38.9745706 |
| Barcelona | Spain | 20 | 41.3850639 | 2.1734035 |

### Whole genome sequencing

Illumina sequencing paired-end libraries were prepared using a custom PCR-Free pipeline including application of dual index primers. Full details of the protocol were described previously in Crawford et al. [13]. Briefly, For samples shipped as carcasses, DNA was extracted using the Chemagic360 DNA tissue extraction kit (PerkinElmer, CMG-723) on a Chemagic360 nucleic acid extractor (PerkinElmer) with a 96-well rod head after samples were homogenized using a steel ball in lysis buffer. Extracted DNA was quantified using Quant-iT PicoGreen (Thermo Fisher Scientific, Waltham, MA, USA), fragmented and double-side size selected to a target size of 350-400 base pairs. Custom barcoded adapters were ligated to fragments, purified, quantified, and pooled for sequencing. Libraries were sequenced using the Illumina HiSeq 4000 or HiSeq X platform producing 151 bp paired-end reads. All fastq files were submitted to the NCBI SRA database under project ID XXXXXXX.

### Short-read processing and quality control

Short-reads were aligned to the AalbF3 (Boyle et al. 2021) *Aedes albopictus* reference sequence (Genbank accession JAFDOQ010000000) using BWA *mem* (https://github.com/lh3/bwa; version 0.7.17-r1188). Duplicate reads were marked using PicardTools (https://broadinstitute.github.io/picard/; version 2.18.10) and bams sorted and filtered using Samtools (http://www.htslib.org/download/; version 0.1.19-96b5f2294a). Since DNA quality was lower in some samples where some DNA fragments were shorter than the target length, we processed all bam files using the clipOverlap program in the bamUtil package (github.com/statgen/bamUtil) to identify and clip paired reads with overlapping ends. We then used GATK (https://github.com/broadinstitute/gatk; version 3.8-1-0-gf15c1c3ef) to realign reads around indels for each population separately.

### Analysis position identification

To minimize effects of sequencing errors and short-read alignment errors, we made VCFs for each population separately using samtools mpileup and the following flags: ‘-q 10 -Q 20 -I -u -t SP,DP’. Putative variant sites were called using BCFtools (1.6, https://samtools.github.io/bcftools/) with ‘-f GQ -c’ flags, which were then filtered using SNPcleaner.pl from ngsQC (https://github.com/tplinderoth/ngsQC) with the following settings ‘-k $MIN_IND -u 2 -D $MAXD -a 0 -f 1e-4 -H 1e-6 -L 1e-6 -b 1e-10’. Sites that passed all filters were retained for further analysis.

### Admixture analysis

We conducted admixture analysis using NGSadmix (). Beagle-style input files for NGSadmix included LD-pruned genotype likelihoods. First, beagle files were made using ANGSD (https://github.com/ANGSD/angsd), with the following command: angsd -bam $BAMS -checkBamHeaders 0 -ref $REF -sites $SITES -out $OUT -minMapQ 20 -minQ 30 -remove_bads 1 -uniqueOnly 0 -GL 1 -doGlf 2 -domajorminor 1. The SITES file included robust sites identified as described above, but excluding sites in the region surrounding the sex locus on chromosome one. To obtain a subset of genomic positions pruned for linkage-disequilibrium, we also prepared input files for PLINK in ANGSD using the following command: angsd -bam $BAMS -checkBamHeaders 0 -ref $REF -sites $SITES -out $OUT -minMapQ 20 -minQ 30 -remove_bads 1 -uniqueOnly 0 --ignore-RG 0 -doPlink 2 -doGeno -4 -doPost 1 -doMajorMinor 1 -GL 1 -doCounts 1 -doMaf 2 -postCutoff 0.9 -geno_minDepth 4 -doGlf 2 -doDepth 1. Output files were used for PLINK LD pruning with the following command: plink --keep-allele-order –tfile $TPED --indep-pairwise 100 10 0.1 --out $OUT --noweb --allow-no-sex --maf 0.01 --geno 0.5 --mind 1. LD-pruned sites were then randomly downsampled to obtain ∼1e6 sites across the genome. Beagle files from above were filtered down to the LD-pruned sites set and used as input for NGSadmix using the following command: NGSadmix -likes $BGL -K $K -P 24 -minInd 1100 -minMaf 0.01 -o $OUT, where $K corresponds to the number of ancestral populations in the model. We modeled K= 2-8.

## Results

We generated over 254 billion paired-end DNA sequence reads from 1,138 individual mosquitoes sampled from 59 localities throughout the global distribution of *Ae. albopictus*. Most regions where this species are found are broadly represented in our dataset including both the native range in South East Asia as well as regions where it is invasive including North East Asia, the Americas, Europe, and Africa (Figure 1).

**Figure 1:**
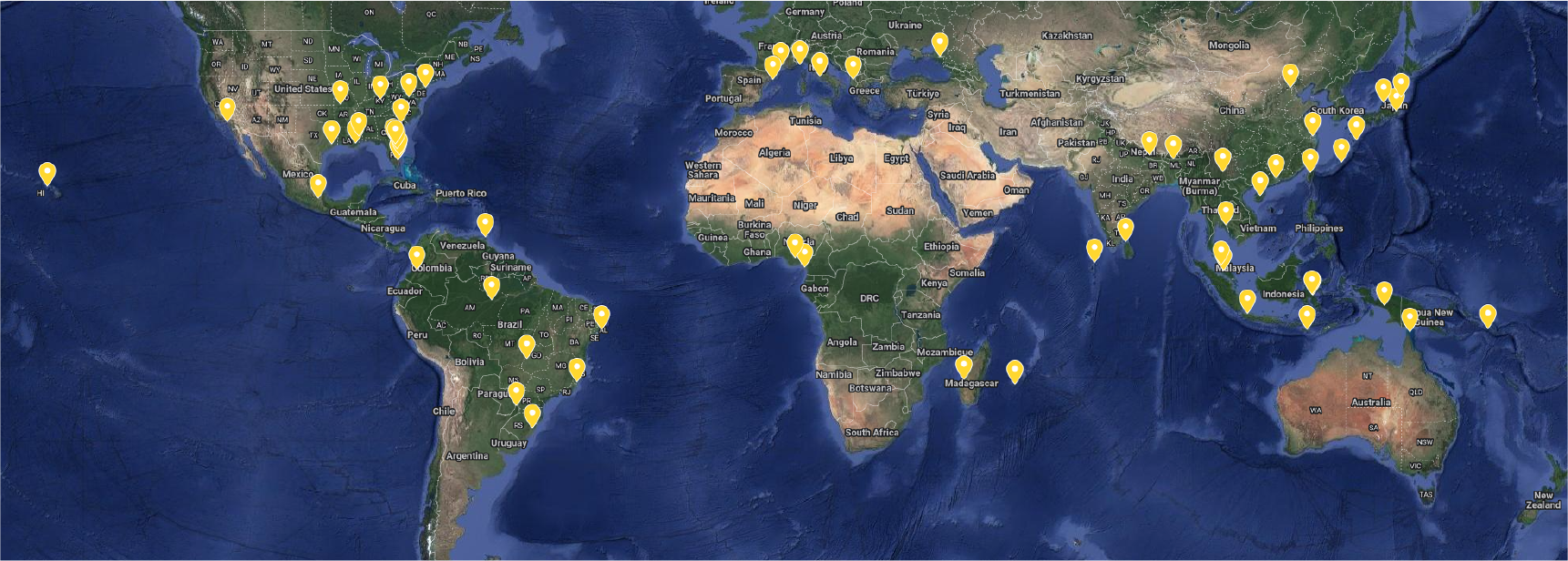
Map of mosquito samples. Yellow markers show location of populations sampled. Samples were included from throughout the distribution of *Ae. albopictus*. See Table 1 for population names and sample sizes.

**Figure 2:**
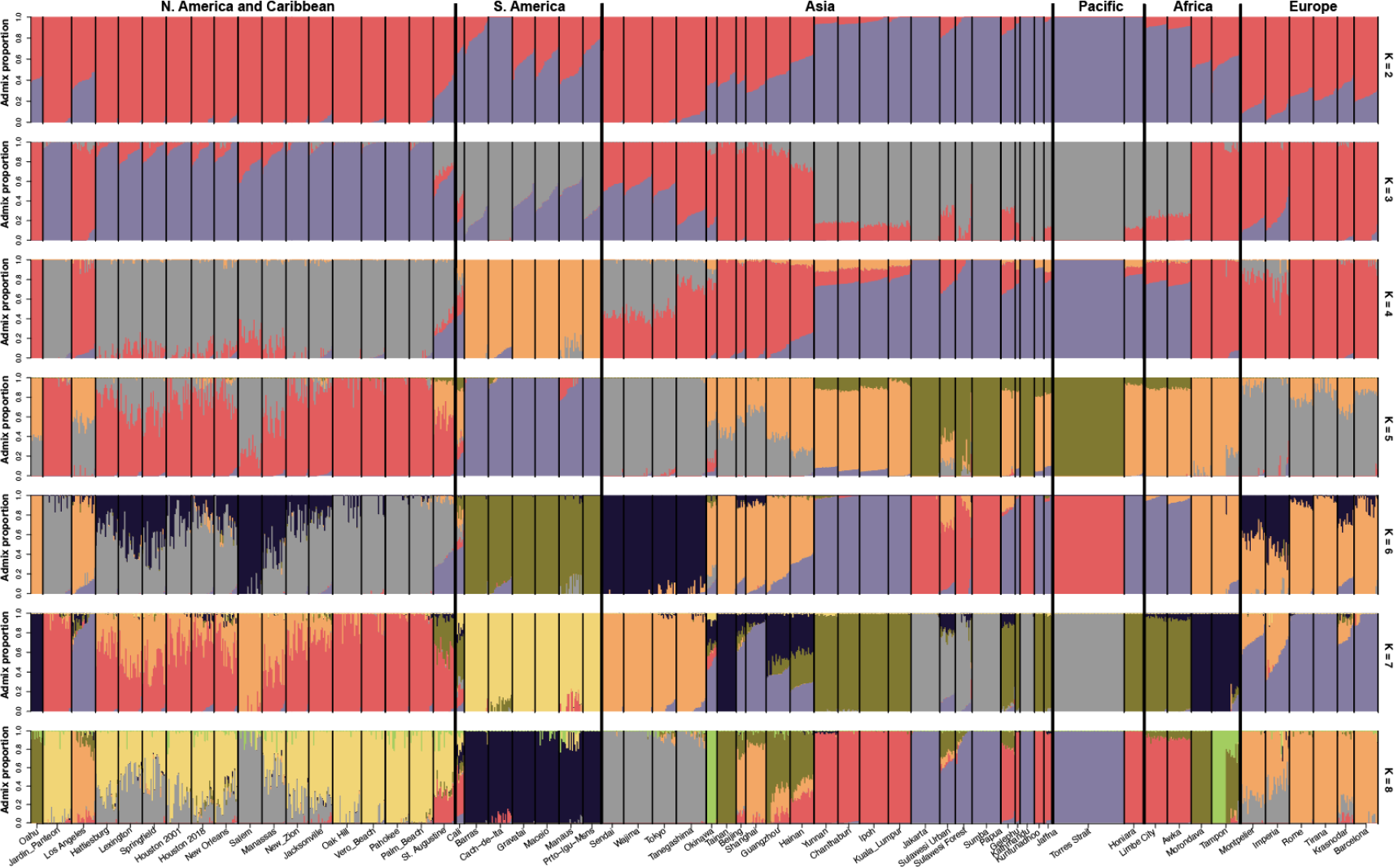
Genetic differentiation and admixture. Barplots show estimates of admixture proportions using NGSadmix with the number of ancestral populations (K) ranging from 2 (top) to 8 (bottom). Each vertical line is one mosquito. Broad regions and continents are delineated by black lines and labels above the plot. Population labels are included below.

We identified high level genetic differentiation in our dataset using admixture analysis and the strongest level of differentiation between populations from the native range in South East Asia and Pacific and those from the invasive range, except for populations from South America that are closer to ancestral populations than to other invasive populations. Increasing the number of ancestral populations (K) to 3 results in North American populations, except for Los Angeles, being assigned to a separate admixture component with large proportions of ancestry in individuals from Japan, Brazil, and Argentina also sharing this component. Increasing the number of ancestral populations reveals regional and continent-level differentiation with several exceptions. For example, at K=8, we see genetic similarity between Oahu (Hawaii), Taiwan, Southern China, and Madagascar, suggesting possible introductions from Southern China to these islands. We also see an interesting component that includes Beijing, Shanghai, Los Angeles, and all of the European countries sampled here. Many populations in the United States show high levels of genetic ancestry shared with populations in Japan.

## Discussion

We describe whole genome sequencing data from 1,125 individual *Ae. albopictus* mosquitoes from 59 localities around the world. These data will be useful to address a number of questions for this species. For example, these data will help understand how this species moves around the world and how it has adapted as it invades new regions and habitats. In particular, photoperiodic diapause has been a transformative adaptation for this species and these data will help to understand the genetic underpinnings of this complex phenotype. These data will also provide a thorough accounting of genetic diversity in this species allowing us to understand how diversity varies across the genome as well as among populations. Work to address many of these questions is ongoing and will be described in forthcoming publications.

In this work, we take a first look at genetic differentiation among the populations sampled here. The high level patterns of differentiation are consistent with those from previous work, with clear differentiation between the tropical and temperate, invasive forms of this species and continent level differentiation generally intact. Several patterns of shared ancestry point to possible sources of introductions, but more work is needed to firm up these findings.

